# Ensemble encoding of facial expressions is greater for static faces than dynamic faces

**DOI:** 10.64898/2025.12.04.692417

**Authors:** Mia Hanson, David Pitcher

**Affiliations:** Department of Psychology, University of York, Heslington, York, YO10 5DD, U.K.

**Keywords:** Facial Expression Discrimination, Ensemble Encoding, Static vs Dynamic Stimuli

## Abstract

Understanding the behaviour and emotions of other people is predicated on accurately decoding nonverbal facial cues. In the current study, we investigated whether the ensemble encoding of these facial cues is processed differently between moving and static faces. Across two experiments, participants performed a delayed-match-to-sample expression recognition task using dynamic and static stimuli. Ensemble size was manipulated by increasing the number of faces in the target array. In Experiment 1, we presented one, two, and four target faces. In Experiment 2, we presented one, four, and eight target faces. Results demonstrated that while recognition accuracy for dynamic and static faces was comparable when one target face was presented, task accuracy diverged as the ensemble size increased. Specifically, increasing the number of faces in the target array failed to improve task performance for the dynamic expression stimuli. In addition, the results of Experiment 2 demonstrated that when eight faces were presented in the dynamic condition, performance was impaired for all expressions (disgust, fear, and happy) as well as in the non-emotional air-puff condition. This suggests motion in general disrupted the participants ability to extract an accurate summary statistic as the ensemble size was increased.

## Introduction

Faces convey a broad range of socially relevant visual cues that are used in everyday interaction and communication. These cues can inform decisions about a person’s identity, gender, age, gaze direction, intent, and emotional state. The ability to efficiently and accurately interpret these cues is a cornerstone of successful social navigation, empathy, and interpersonal interaction. Cognitive and neurobiological models of face processing (Allison et al., 2000; Bruce & Young, 1986; Haxby et al., 2000; Pitcher & Ungerleider, 2021) propose that these cues can be broadly segregated into cues that rely on invariant aspects (e.g. facial structure and shape) and changeable aspects (e.g. eye, mouth and head movements). The invariant (or static) cues contribute to identity discrimination (Grill-Spector et al., 2004; Megreya & Burton, 2006; Sliwinska et al., 2022) while the changeable aspects contribute to facial expression discrimination (Krumhuber et al., 2023; Pitcher et al., 2020). This functional dissociation between static and dynamic facial cues is supported by evidence from different methods. These include behavioural experiments (Calvo et al., 2016; Dobs et al., 2018; Lander et al., 1999), neuropsychology (Pitcher, 2025; Prabhakar et al., 2025; Sliwinska, Bearpark, et al., 2020), neuroimaging (Fox et al., 2009; Kucuk et al., 2024; Puce et al., 1998; Schultz & Pilz, 2009) and developmental studies (Kosakowski et al., 2022).

The majority of behavioural research into facial expression discrimination has used static images as experimental stimuli (Dawel et al., 2022). Static stimuli, typically photographs of peak emotional expressions, offer a controlled and precise method for examining the recognition of affective information from discrete, snapshot-like cues (Ekman & Friesen, 1976; Pitcher, 2014; Pitcher et al., 2008; Pitcher et al., 2023; Sliwinska, Elson, et al., 2020). However, in the real world, faces are rarely static. They are constantly in motion as facial expressions unfold over time. This temporal dimension is a rich source of socially relevant information (Fiorentini et al., 2012; Krumhuber & Scherer, 2011; Krumhuber et al., 2023). Dynamic stimuli provide this critical temporal information and reveal the unique trajectory and biomechanics of how an expression is formed and how it dissipates. Consequently, some empirical evidence has reported an advantage for expression recognition using dynamic stimuli (Arsalidou et al., 2011; Fiorentini & Viviani, 2011; Jiang et al., 2014; Kamachi et al., 2001; Richoz et al., 2024). In the current study we investigated whether dynamic stimuli would also produce a behavioural advantage when recognising multiple facial expressions simultaneously, a process known as ensemble encoding.

Ensemble encoding is the cognitive process in which observers are able to extract an average summary statistic from multiple exemplars of a stimulus category rather than processing each exemplar individually (Whitney & Yamanashi Leib, 2018). This produces a redundancy gain, where the coactivation of multiple signals increases the overall signal strength and results in faster, more accurate task performance. Prior research has demonstrated that expression recognition benefits from ensemble encoding when using both static (Haberman & Whitney, 2007, 2009; Hao et al., 2025; Sama et al., 2025) and dynamic (Elias et al., 2017) stimuli. However, it is unclear whether the redundancy gain is the same for both dynamic and static expressions. The additional visual information conveyed in dynamic stimuli may interfere with the ability to form an efficient ensemble summary. Instead of a redundancy gain, processing multiple dynamic faces might result in a redundancy cost, impairing performance as the number of faces increases.

To test this hypothesis, we designed two experiments utilizing a delayed match-to-sample task facial expression discrimination task. Participants were asked to match facial expressions presented as either dynamic videos or as static images of the peak expression (van der Gaag et al., 2007). We systematically manipulated ensemble size by varying the number of faces presented in the target array (Figure 1a). In Experiment 1, we compared accuracy for one, two, and four target faces. In Experiment 2, we compared accuracy for one, four, and eight target faces. Facial expression stimuli included a range of basic emotions (happy, fear, disgust) as well as neutral faces (the actor staring expressionless into the camera) and a non-emotional motion control (“air-puff”) to determine if these any differential effects were limited to facial expressions or generalised to any facial movement. We hypothesised that ensemble encoding may be more challenging when using dynamic compared to static stimuli as the visual information needs to be summated over a longer time scale.

**Figure 1.**
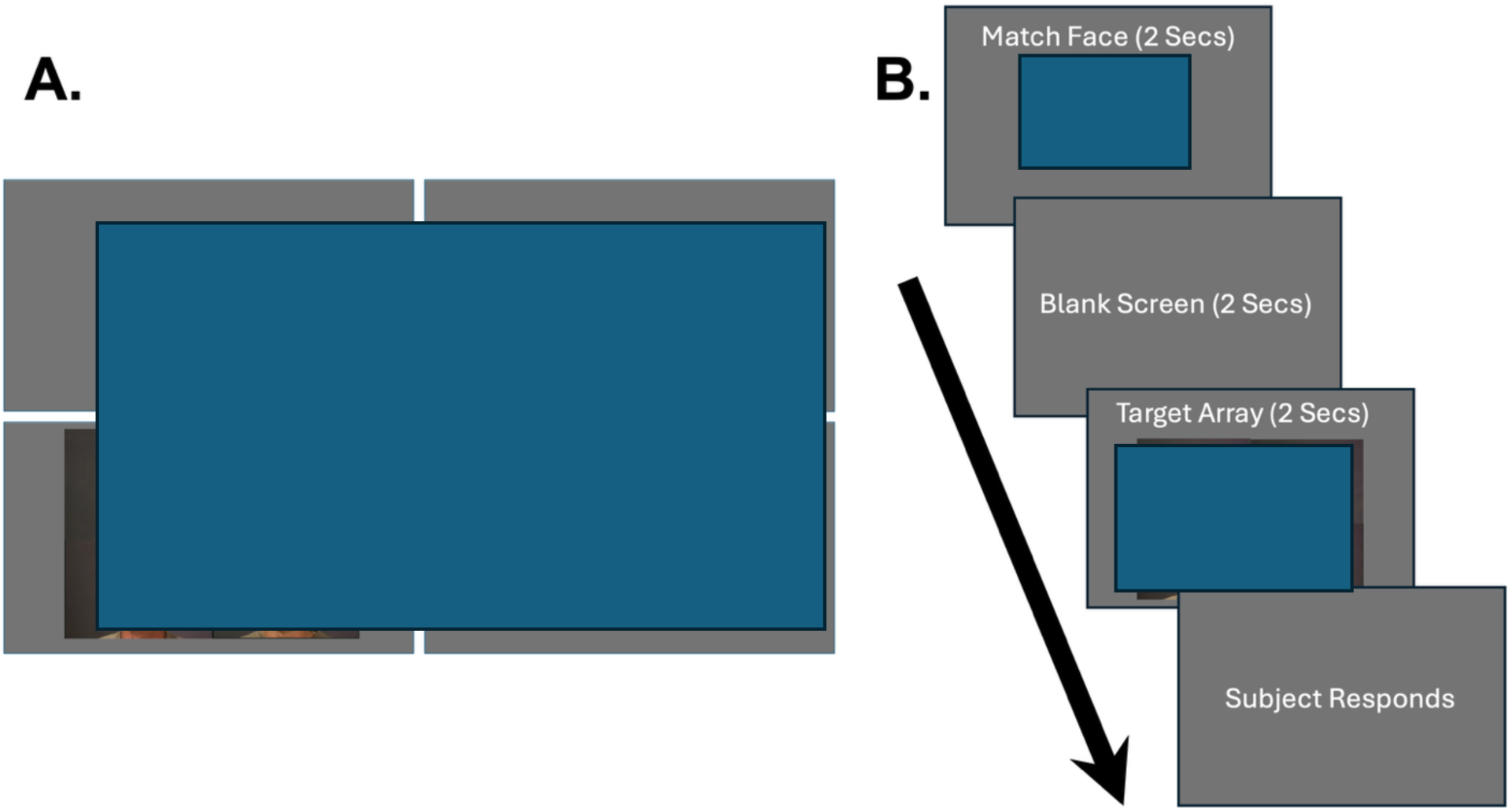
a. Shows the format of the Target Arrays for the static condition. The dynamic arrays presented videos in the same configuration. In Experiment 1 participants saw arrays with 1, 2 or 4 faces. In Experiment 2 participants saw arrays with 1, 4 or 8 faces. The stimuli in the arrays for the 2, 4 and 8 faces all expressed the same emotion. Figure 1b. The trial procedure for Experiment 1 and 2. Participants performed a delayed match to sample expression discrimination task. Participants had to indicate whether the expression of the match face was the same as the faces (or face) in the target array.

## Materials and Methods

### Participants

50 participants (35 females, 13 males and 2 non-binary) took part in Experiment 1. 40 participants (27 females, 9 males and 4 non-binary) took part in Experiment 2. All subjects had normal or corrected-to-normal vision and gave informed consent as directed by the ethics committee in the Psychology department at the University of York.

### Stimuli

Dynamic stimuli consisted of videos of actors depicting different emotional facial expressions (van der Gaag et al., 2007). These stimuli have been used in previous studies of emotion discrimination (Nikel et al., 2022; Pitcher et al., 2020). The videos show one of twelve actors (6 males, 6 females, all Caucasian) depicting different facial expressions (disgust, fear, happy) together with the actors blowing out cheeks (motion with no expression) and neutral (in which the actors stare into the camera without expression). This resulted in a total of 60 video stimuli. Static stimuli were created using a single frame of peak emotional expression from each video. All stimuli were then organised by emotion and numbered from 1-12 based on the actor (e.g. happy 1, disgust 5, etc.). Stimuli were presented in full colour on a grey background. Dynamic stimuli were presented without audio. Dynamic and static stimuli were shown for 2 seconds at a frame rate of 143 frames per second.

### Procedure

Participants performed a delayed-match-to-sample facial expression discrimination task using dynamic and static stimuli (Figure 1a). The experiment was conducted using Psychopy Version 3 (Peirce et al., 2019), which controlled randomisation and presentation of the stimuli, as well as measuring response time and accuracy. Each trial began with a single match face (dynamic or static) presented centrally for 2 seconds. This was followed a blank grey screen for two seconds. The target face array (dynamic or static) was then presented. In Experiment 1 the target array consisted of 1 face (Figure 1b), 2 faces (Figure 1c) or 4 faces (Figure 1d). In Experiment 2 the target array consisted of 1 face (Figure 1b), 4 faces (Figure 1d) or 8 faces (Figure 1e). All faces in the target arrays presented the same expressions in all conditions (e.g. all 8 videos in the dynamic condition would all depict the same expression). The participants were asked to match if the expression of the match face was the same as the face, or faces, depicted in the target arrays. This was done using the arrow keys to indicate whether faces in the target array showed the same emotion (right arrow key) or a different emotion (left arrow key) to the match face. In Experiment 1 participants completed six blocks: dynamic stimuli with 1, 2 and 4 face arrays and static stimuli with 1, 2 and 4 face arrays. Each block consisted of fifty trials. In Experiment 2 participants completed six blocks: dynamic stimuli with 1, 4 and 8 face arrays and static stimuli with 1, 4 and 8 face arrays. Each block consisted of fifty trials. Block order was randomised across participants.

## Results

### Experiment 1

The results of Experiment 1 (Figure 2a) demonstrated that static facial expressions were more accurately matched than dynamic facial expressions but only when two or four target faces were presented. Accuracy data were entered into a two (motion: dynamic and static) by three (number of faces: one face, two faces and four faces) by five (emotion: happy, neutral, air-puff, fear and disgust) repeated measures analysis of variance (ANOVA). We found significant main effects of motion (F (1,49) = 15.44, *p* <0.001; partial η ^2^ = 0.24), number of faces (F (2, 98) = 5.48, *p* = 0.006; partial η ^2^ = 0.10) and emotion (F (4,196) = 37.65, *p* = 0.001; partial η ^2^ = 0.44). Motion and number of faces (F (2,98) = 6.01, *p* = 0.003; partial η ^2^ = 0.11) and motion and emotion (F (4,196) = 20.50, *p* < 0.001; partial η ^2^ = 0.30) both combined in significant two-way interactions. Number of faces and emotion failed to show a significant two-way interaction (F (8,392) = 0.94, *p* = 0.48; partial η ^2^ = 0.02). There was no significant three-way interaction between motion, number of faces and emotion (F (8,392) = 0.85, *p* = 0.56; partial η ^2^ = 0.02).

**Figure 2.**
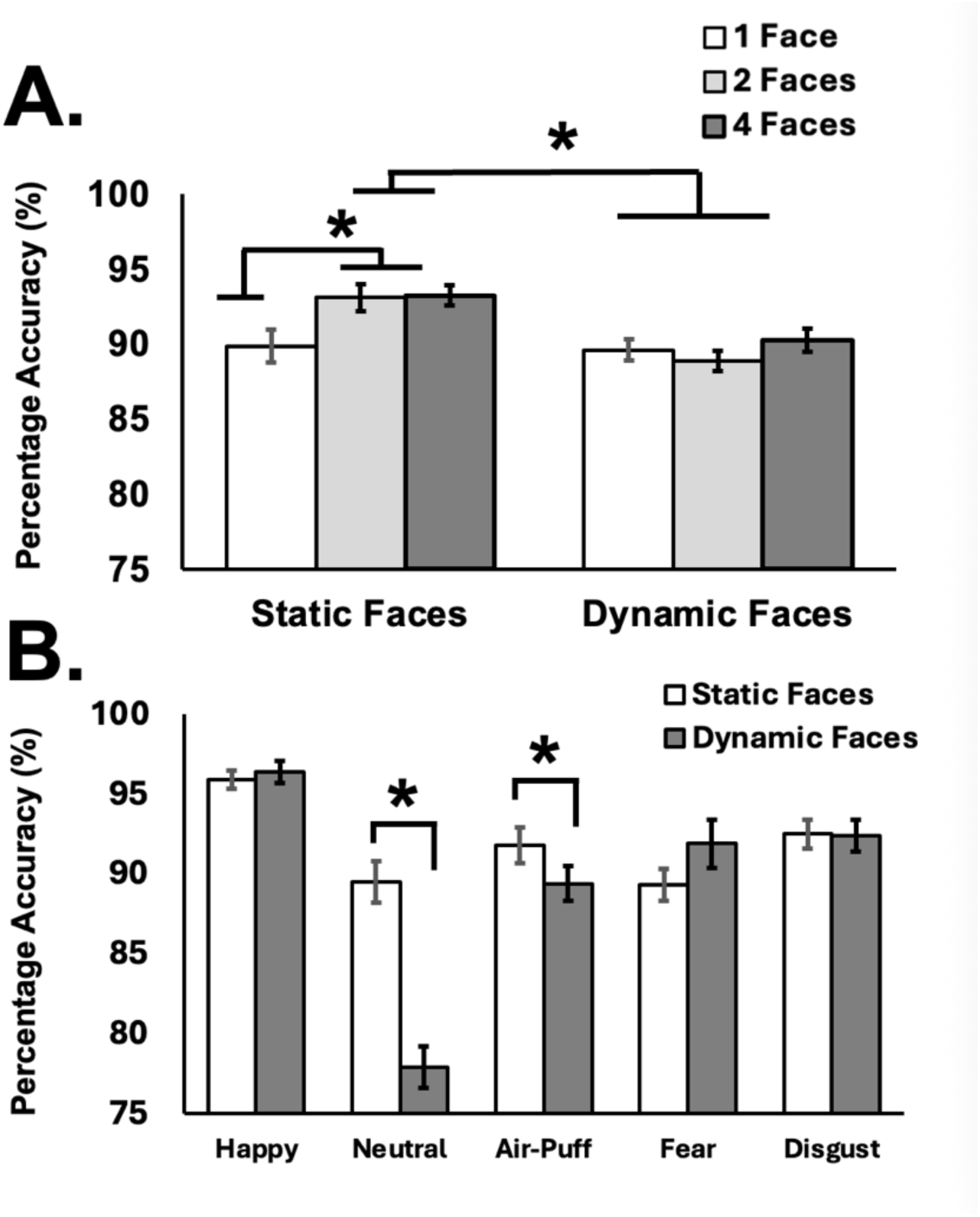
a. The results of the significant two-way ANOVA between motion and number of faces in the target array. Pairwise comparisons showed that static faces were more accurately recognized than dynamic faces when 2 faces (*p* < 0.001) and 4 faces (*p* < 0.001) were presented. Figure 2b. The results of the significant two-way ANOVA between motion and emotion. Pairwise comparisons showed significant differences between dynamic and static faces when the neutral (*p* < 0.001) and air-puff faces (*p* = 0.044). were presented

To understand what factors were driving the significant two-way interaction between motion and number of faces we analysed the data in a separate ANOVA (Figure 2a). Results showed significant main effects of motion (F (1,49) = 19.25, *p* < 0.001; partial η ^2^ = 0.28), number of faces (F (2,98) = 5.00, *p* = 0.009; partial η ^2^ = 0.09) and a significant two-way interaction (F (2,98) = 4.57, *p* = 0.013; partial η ^2^ = 0.85). Pairwise comparisons showed that static faces were more accurately recognized than dynamic faces when 2 faces (*p* < 0.001) and 4 faces (*p* < 0.001) were presented. There was no significant difference between dynamic and static faces when 1 face was presented (*p* = 0.82). Further comparisons showed significant differences in the static condition between the 1 and 2 target face condition (*p* = 0.027) and between the 1 and 4 target face condition (*p* = 0.008) but not between the 2 and 4 target face condition (*p* = 1). There were no significant differences in the dynamic face condition between any of the target face conditions (*p* > 0.26).

Next, to understand what factors were driving the significant two-way interaction between motion and emotion we analysed the data in a separate two-way ANOVA (Figure 2b). Results showed significant main effects of motion (F (1,49) = 15.44, *p* = 0.001; partial η ^2^ = 0.24), emotion (F (4,196) = 37.65, *p* < 0.001; partial η ^2^ = 0.44) and a significant two-way interaction (F (4,196) = 20.50, *p* < 0.001; partial η ^2^ = 0.30). Pairwise comparisons showed significant differences between dynamic and static faces when the neutral expressions were presented (*p* < 0.001) and for air-puff expressions (p = 0.044), but not when happy (*p* = 0.5), fear (*p* = 0.09) or disgust (*p* = 0.8) expressions were presented.

Reaction time (RT) data were entered into a two (motion: dynamic and static) by three (number of faces: one face, two faces and four faces) by five (emotion: happy, neutral, air-puff, fear and disgust) repeated measures analysis of variance (ANOVA). We found significant main effects of motion (F (1,49) = 280.00, *p* < 0.001; partial η ^2^ = 0.85), number of faces (F (2,98) = 8.04, *p* < 0.001; partial η ^2^ = 0.14) and emotion (F (4,196) = 18.30, *p* < 0.001; partial η ^2^ = 0.27). Motion and emotion combined in significant two-way interaction (F (4,196) = 3.96, *p* = 0.004; partial η ^2^ = 0.08). However, there were no two-way interactions between motion and number of faces (F (2,98) = 0.15, *p* = 0.86; partial η ^2^ = 0.003) or between emotion and number of faces (F (8,392) = 0.6, *p* = 0.8; partial η ^2^ = 0.012). There was no significant three-way interaction between motion, number of faces and emotion (F (8,392) = 0.58, *p* = 0.79.; partial η ^2^ = 0.012). Pairwise comparisons of the significant main effects showed that static faces were more quickly recognised than dynamic faces (*p* < 0.001), that 4 faces in the target array were more quickly recognised than 1 face (*p* = 0.001) and happy expressions were more quickly recognised than all other expressions (*p* < 0.001). No other tests reached significance (*p* > 0.075).

### Experiment 2

Accuracy data (Figure 3) were entered into a two (motion: dynamic and static) by three (number of faces: one face, four faces and eight faces) by five (emotion: happy, neutral, air-puff, fear and disgust) repeated measures analysis of variance (ANOVA). We found significant main effects of motion (F (1,39) = 43.37, *p* < 0.001; partial η ^2^ = 0.53), number of faces (F (2,78) = 7.89, *p* < 0.001; partial η ^2^ = 0.17) and emotion (F (4,156) = 18.41, *p* < 0.001; partial η ^2^ = 0.32). Motion and number of faces (F (2,78) = 6.14, *p* = 0.003; partial η ^2^ = 0.14), motion and emotion (F (4,156) = 6.02, *p* < 0.001; partial η ^2^ = 0.13) and number of faces and emotion (F (8,312) = 2.06, *p* = 0.039; partial η ^2^ = 0.05). Finally, motion, number of faces and emotion combined in a significant three-way interaction (F (8,312) = 4.51, *p* < 0.001; partial η ^2^ = 0.10).

**Figure 3.**
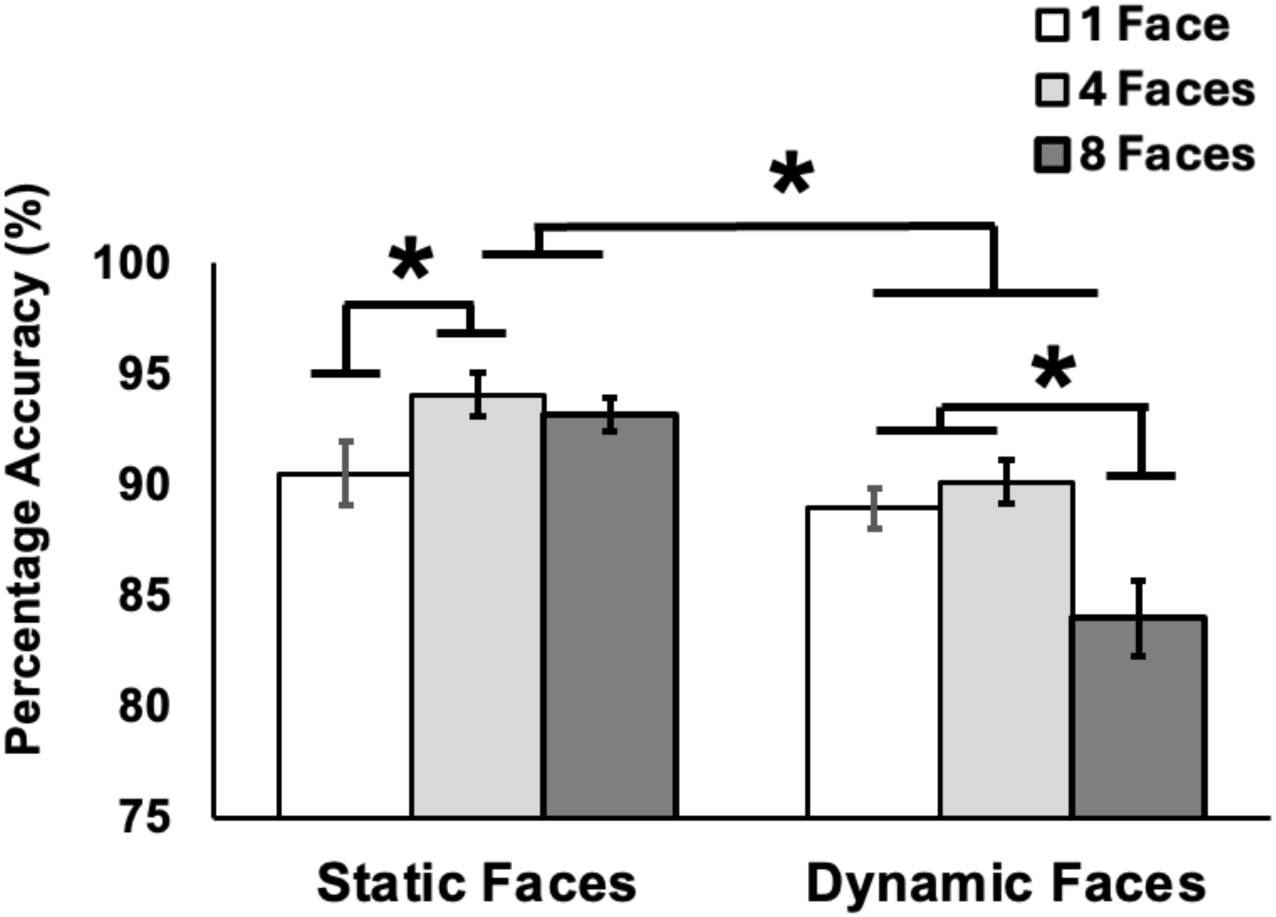
The results of the significant two-way ANOVA between motion and number of faces in the target array. Pairwise comparisons showed that static faces were more accurately recognized than dynamic faces when 4 faces (p < 0.001) and 8 faces (p < 0.001) were presented. For dynamic faces there was a no significant difference between when 1 and 4 faces (p = 0.026) and when 1 and 8 (p = 0.003) faces were presented.

To understand what factors were driving the significant three-way interaction we analysed the data in separate two-way ANOVAs. Firstly, we ran an ANOVA between motion and number of faces to compare the results of experiment 2 (in which 1, 4 and 8 target faces were presented) with experiment 1 (in which 1, 2 and 4 target faces were presented).

Results (Figure 3) showed significant main effects of motion (F (1,39) = 43.64 , *p* < 0.001; partial η ^2^ = 0.53), number of faces (F (2,78) = 7.74, *p* < 0.001; partial η ^2^ = 0.17) and a significant two-way interaction (F (2,78) = 7.42, *p* = 0.001; partial η ^2^ = 0.16). Pairwise comparisons showed that static faces were more accurately recognized than dynamic faces when 4 faces (*p* < 0.001) and 8 faces (*p* < 0.001) were presented. There was no significant difference between dynamic and static faces when 1 face was presented (*p* = 0.29). Further comparisons showed significant differences in the static condition between the 1 target face and 4 target faces condition (*p* = 0.02) but not between the 1 target face and 8 target faces condition (*p* = 0.11) or between the 4 and 8 target face condition (*p* = 0.98). In the dynamic condition there was no significant difference between the 1 and 4 target faces (*p* = 0.86) but there were between when 1 and 4 faces (*p* = 0.03) and when 1 and 8 (*p* = 0.003) faces were presented.

Next, to examine whether there were significant differences between the discrimination of dynamic and static facial expression across the target face conditions we performed separate ANOVAs comparing dynamic and static stimuli with expressions for each of the target array conditions (Figure 4).

**Figure 4.**
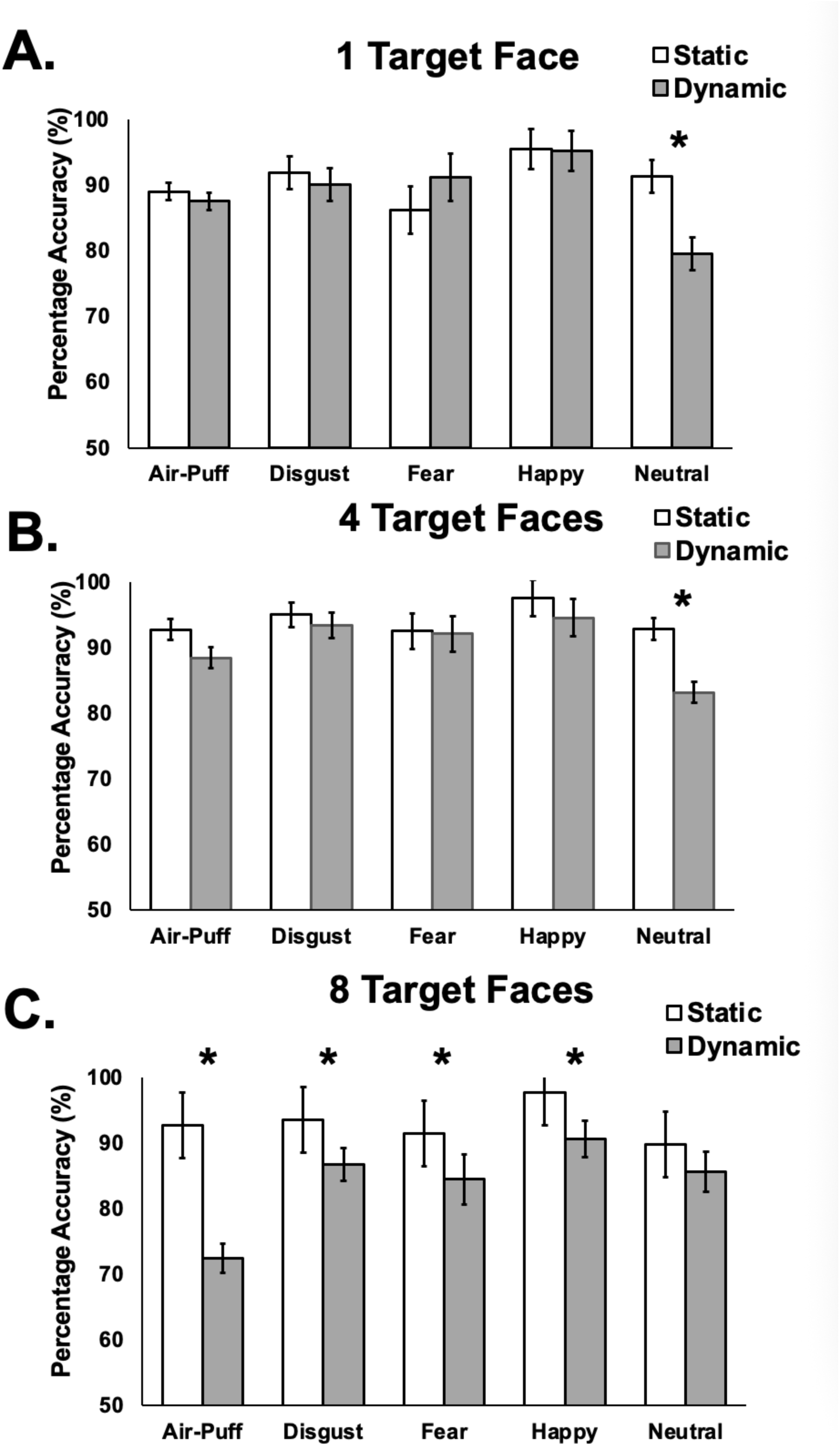
a. The results of the significant ANOVA between motion conditions (dynamic or static) and expression for when 1 target face was presented. Pairwise comparisons showed a significant difference for neutral expressions only (p < 0.001). Figure 4b. The results when 4 target faces were presented. The only significant difference was in the neutral face condition (p < 0.001). Figure 4c. The results when 8 target faces were presented. There were significant differences between dynamic and static when air-puff (p < 0.001), disgust (p = 0.033), fear (p = 0.017) and happy (p = 0.003 expressions were presented but not when neutral expressions were presented (p = 0.1).

<u>1 Target Face</u>: Accuracy data when 1 face was presented were analysed in a two-way ANOVA (Figure 4a). Results did not show a significant main effect of motion (F (1, 39) = 1.87, *p* = 0.18; partial η ^2^ = 0.46). There was a significant main effect of expression (F (4,156) = 10.28, *p* < 0.001; partial η ^2^ = 0.21) and a significant two-way interaction (F (4,156) = 5.80, *p* < 0.001; partial η ^2^ = 0.13). Pairwise comparisons showed significant differences between dynamic and static faces when the neutral expressions were presented (*p* < 0.001) but not when happy (*p* = 0.81). air-puff (*p* = 0.68), fear (*p* = 0.11) or disgust (*p* = 0.47) expressions were presented.

<u>4 Target Faces</u>: Accuracy data when 4 faces were presented were analysed in a two-way ANOVA (Figure 4b). Results showed significant main effects of motion (F (1, 39) = 20.99, *p* < 0.001; partial η ^2^ = 0.35) and expression (F (4,156) = 6.35, *p* < 0.001; partial η ^2^ = 0.14) and a significant two-way interaction (F (4,156) = 2.62, *p* = 0.037; partial η ^2^ = 0.06). Pairwise comparisons showed significant differences between dynamic and static faces when the neutral expressions were presented (*p* < 0.001) but not when happy (*p* = 0.07), air-puff (*p* = 0.12), fear (*p* = 0.88) or disgust (*p* = 0.31) expressions were presented.

<u>8 Target Faces</u>: Accuracy data when 8 faces were presented were analysed in a two-way ANOVA (Figure 4c). Results showed significant main effects of motion (F (1, 39) = 26.56, *p* < 0.001; partial η ^2^ = 0.41) and expression (F (4,156) = 9.40, *p* < 0.001; partial η ^2^ = 0.19) and a significant two-way interaction (F (4,156) = 6.04, *p* < 0.001; partial η ^2^ = 0.13). Pairwise comparisons showed significant differences between dynamic and static faces when happy (*p* = 0.003), air-puff (*p* < 0.001), fear (*p* = 0.016) and disgust (*p* = 0.03) expressions were presented but not when neutral expressions were presented (*p* = 0.10).

Reaction time (RT) data were entered into a two (motion: dynamic and static) by three (number of faces: one face, two faces and four faces) by five (emotion: happy, neutral, air-puff, fear and disgust) repeated measures analysis of variance (ANOVA). We found significant main effects of motion (F (1,39) = 653.18, *p* < 0.001; partial η ^2^ = 0.94), number of faces (F (2,78) = 10.62, *p* < 0.001; partial η ^2^ = 0.21) and emotion (F (4,156) = 9.96, *p* = 0.001; partial η ^2^ = 0.20). Both motion and emotion (F (4,156) = 5.73, *p* < 0.001; partial η ^2^ = 0.13) and number of faces and emotion F (8,312) = 2.31, *p* = 0.02; partial η ^2^ = 0.06) combined in significant two-way interactions. There was no two-way interaction between motion and number of faces F (2,78) = 0.34, *p* =0.72; partial η ^2^ = 0.009). There was no significant three-way interaction between motion, number of faces and emotion (F (8,312) = 1.42, *p* = 0.19.; partial η ^2^ = 0.035). Pairwise comparisons of the significant main effects showed that static faces were more quickly recognised than dynamic faces (*p* < 0.001). 8 faces were more quickly recognised than 4 faces (*p* = 0.028 and 1 face (*p* < 0.001). Happy expressions were more quickly recognised than all other expressions (*p* < 0.004). No other tests reached significance (*p* > 0.5).

## Discussion

In the current study we investigated whether ensemble encoding of facial expressions would differ when using dynamic and static stimuli. We hypothesised that the redundancy gain reported for static facial expression recognition (Haberman & Whitney, 2007, 2009; Hao et al., 2025; Sama et al., 2025) may not be as robust when using dynamic facial expression stimuli. Our results support this hypothesis. Across two experiments, we demonstrated that while discrimination accuracy for dynamic and static faces was comparable when viewing a single target face, a performance difference emerged as the number of faces in the target array was increased. Specifically, static expressions were recognized significantly more accurately than dynamic expressions when the target array contained two or four faces (Figure 2a) and when the target array contained four or eight faces (Figure 3). Furthermore, Experiment 2 demonstrated that for dynamic stimuli, a target array of eight target faces resulted in significantly worse performance than target arrays containing four faces or one face. This impaired performance for dynamic stimuli was observed for all three expressions (disgust, fear and happy) as well as in the non-emotional motion “air-puff” condition (Figure 4c). This suggests that facial motion in general, rather than motion associated only with facial expressions, disrupts efficient ensemble encoding.

The difference in performance between dynamic and static stimuli was driven by the increase in performance accuracy for the static stimuli when the target array contained two or four faces (Figure 2a and Figure 3). This improvement is consistent with theories proposing a redundancy gain in task performance when two (or more) signals are coactivated resulting in faster and more accurate task performance (Miller, 1982). Our results suggest that when two or four faces were presented in the static target arrays the correspondence of the faces expressing the same emotion increased the strength of the signal and improved task performance. In Experiment 2 there was an increase in performance when eight faces were presented in the target array, but the difference was not statistically significant (Figure 3). This suggests that eight static faces may be reaching the limit where the increase in signal strength is effective. Crucially, across both experiments we failed to see this increased accuracy for dynamic stimuli suggesting that dynamic faces result in a redundancy cost rather than a redundancy gain. This cost has been reported in prior studies that required participants to match identity and expression using static stimuli (Yankouskaya et al., 2012; Yankouskaya, Humphreys, et al., 2014; Yankouskaya, Rotshtein, et al., 2014).

Extracting relevant social information from dynamic faces requires processing the temporal unfolding of motion, biomechanics, and trajectory of facial components (Krumhuber et al., 2023). The complex, time-varying information from multiple faces in the target arrays may have been difficult to integrate into a single, stable ensemble statistic. In contrast, processing multiple static images of peak expressions may allow for a more rapid and efficient averaging of features (Haberman & Whitney, 2007, 2009; Hao et al., 2025; Sama et al., 2025). In our single-face condition, this computational cost of processing motion is not aggregated, and resources are sufficient for both tasks. However, when required to form an ensemble using multiple faces, the cognitive demands of integrating multiple dynamic signals outweighs the potential benefit of redundancy, leading to a redundancy cost. This pattern of results is consistent with cognitive and neurobiological models that propose dissociable pathways for the processing of invariant (static) and changeable (dynamic) facial cues (Allison et al., 2000; Bruce & Young, 1986; Haxby et al., 2000; Pitcher, 2021; Pitcher & Ungerleider, 2021). The behavioural dissociation we report suggests that the dynamic processing pathway is more susceptible to disruption when cognition is taxed while the static pathway is more resilient. This suggests that these pathways are not just functionally but are also computationally distinct.

Our findings help contextualize the dynamic advantage reported in previous studies of facial expression discrimination (Arsalidou et al., 2011; Fiorentini & Viviani, 2011; Jiang et al., 2014; Kamachi et al., 2001; Richoz et al., 2024). This advantage, typically investigated using single faces, may not scale to group or ensemble perception. Our study demonstrates that any benefit of dynamic information for a single face is lost when multiple faces must be processed concurrently. Interestingly a previous study using dynamic faces did report an ensemble advantage for facial expression recognition (Elias et al., 2017). This study did not compare dynamic and static face stimuli but instead contrasted synchronous and asynchronous expressions using dynamic stimuli. In addition, the faces were originally static stimuli (Tottenham et al., 2009) and the dynamic aspect was created by morphing between expressions. These differences may account for the discrepancy between our studies but further experiments will be needed to address this discrepancy.

We argue that the informational richness of dynamic cues comes at a computational cost. In real-world conditions, such as trying to track the expressions of multiple people in a crowded room, our data suggest that the complexity of integrating multiple dynamic motion signals becomes a liability. Efficiently summarizing the “average mood” of a crowd may paradoxically rely more on processing static, peak expression cues than on the full, complex dynamic information. This interpretation is bolstered by the results for the neutral faces. In Experiment 1, the static advantage was driven by the neutral faces only (Figure 2b). This pattern also appeared in Experiment 2 in the one and four face target array conditions (Figure 4a and 4b). This was likely due to the nature of the stimuli. The dynamic neutral faces (an actor staring into the camera) may have been perceived as ambiguous, with subtle, non-expressive movements (e.g., blinks) being misinterpreted as suppressed intent or emotion. However, when the eight face target array was shown, this pattern reversed (Figure 4c). The difference for neutral faces was no longer significant, while a significant static face advantage emerged for all three emotional expressions (disgust, fear, and happy) and for the non-emotional motion control (air-puff). This suggests two distinct mechanisms. At low-to-moderate ensemble sizes, the dominant effect was the ambiguity of dynamic-neutral stimuli. At the highest ensemble size, this ambiguity effect was less disruptive than the general computational cost of processing complex facial motion from eight distinct sources, which impaired performance across all dynamic conditions.

Our study provides novel behavioural evidence that forming an ensemble representation of dynamic facial expressions is a computationally demanding process. These findings show the crucial trade-off between the informational richness of dynamic cues and their cognitive computational cost when processed as an ensemble. Real world social situations will sometimes require the processing of many different facial expressions. Our results demonstrate that this is a computationally challenging process and future studies could manipulate real-world social interactions to better understand these processes (Sliwinska et al., 2022). Furthermore, this paradigm could be a powerful tool for investigating atypical social perception in clinical populations known to have difficulties with facial expression recognition.

## Acknowledgements

This work was supported by Leverhulme Trust Project Grant RPG-2024-389 (D.P.) and a summer student scholarship from the Department of Psychology at the University of York (M.H.). The authors thank Mike Burton for advice on experimental design.

## Statements and declarations

### Ethical considerations

The study was approved by the ethics committee in the Psychology department at the University of York.

### Consent to participate

All participants gave informed consent

### Consent for publication

All images have received consent for publication.

